# Navigating the Valley of Death: Perceptions of Industry and Academia on Production Platforms and Opportunities

**DOI:** 10.1101/2020.05.04.075770

**Authors:** Linde FC Kampers, Enrique Asin Garcia, Peter J Schaap, Annemarie Wagemakers, Vitor AP Martins dos Santos

## Abstract

Rational lifestyle engineering using computational methods and synthetic biology has made it possible to genetically improve industrial performance of microbial cell factories for the production of a range of biobased chemicals. However, only an estimated 1 in 5,000 to 10,000 innovations make it through the Valley of Death to market implementation.

To gain in-depth insights into the views of industry and academia on key bottlenecks and opportunities to reach market implementation, a qualitative and exploratory study was performed by conducting 12 in depth interviews with 8 industrial and 4 academic participants. The characteristics that any cell factory must have were schematically listed, and commonly recognised opportunities were identified.

We found that academics are limited by only technical factors in their research, while industry is restricted in their research choices and flexibility by a series of technical, sector dependent and social factors. This leads to a misalignment of interest of academics and funding industrial partners, often resulting in miscommunication. Although both are of the opinion that academia must perform curiosity-driven research to find innovative solutions, there is a certain pressure to aim for short-term industrial applications. All these factors add up to the Valley of Death; the gap between development and market implementation.

A third party, in the form of start-up companies, could be the answer to bridging the Valley of Death.

## Introduction

The demand for industrial production by micro-organisms is ever increasing. In 1990, the global market value of industrial enzymes was close to a billion USD, crossed the two billion USD mark in 2005 [1], was valued at over four and a half billion USD in 2016 and is expected to reach over six billion USD in 2022 [2]. Nowadays, bacteria, yeasts, fungi, and micro-algae are used in the industrial production processes of food, enzymes, vitamins, pharmaceuticals, biofuels, bioplastics, bio-insecticides, nanocomposites for electronic devices and a large variety of chemicals and enzymes with industrial value. The reasons to widely apply a diverse set of micro-organisms are manifold; they represent a broad biochemical diversity, increase the feasibility of large scale production, can produce sustainable products, reduce processing time, require low energy input, increase cost effectiveness and can be selected for non-toxicity [3, 4, 5, 6, 7, 8, 9, 10, 11, 12].

However, as heavily as the current biotechnology industry depends on production by microbes, production methods are by far not as stable as desired due to the biological variability of micro-organisms [13]. Many external factors affect the production capacity of microorganisms, including temperature, pH, oxygen, ion and carbon source availability, cell density, and biofilm formation [14, 15, 16, 17, 18, 19, 20, 21, 22, 23]. All factors combined, this often results in a high variability in yield or product profile.

For industrial production purposes it might be ideal to select only one microorganism as the general production platform. By focusing research on a single strain, the microbial toolkit would increase, production would stabilize, and production costs would decrease as more universal bioreactor set-ups could be used. An ideal industrial microbial workhorse must be safe to work with (preferably generally regarded as safe, or GRAS), genomically accessible, metabolically and environmentally flexible, resistant to external industrial stress factors, grow on cheap medium, and capable to produce a high variety of products with high and stable yields ‘a la carte (upon appropriate tailoring). The field of systems and synthetic biology potentially offers rational solutions to these requirements. By restructuring the genome and thus the metabolism of micro-organisms, microbes found in nature can be perfected to produce any ‘a la carte product of interest in a safe way. Whereas random evolution and selection are common methods to increase productivity, yield or performance of a species, in silico systems and in vivo synthetic biology offers the tools to not only adapt microorganisms to gain novel characteristics in a directed way, but to create them. This goes far beyond the engineering of metabolic pathways for production of compounds, and extends to re-programming the lifestyle of microbes to operate outside of their natural boundaries.

An example of research conducted in academia to improve microbial workhorses for robust biocatalysis is one reported by Sandberg and colleagues, who aimed to render the industrially-relevant Escherichia coli more resilient to temperature fluctuations [25]. In this study, E. coli K-12 MG1655, which has an optimal growth temperature at 37◦C, was subjected to adaptive laboratory evolution to improve strain performance at 42◦C. Another example is the lifestyle adaptation of the industrially applied Pseudomonas putida, which was enriched with three heterologous genes to better survive micro-oxic conditions through oxygen gradients [19]. So far, however, these examples are not applied in industry.

The large gap for academic research to advance from the technological demonstration to actual commercialization in industry is generally referred to as the “Valley of Death” [26]. In biotechnology, including pharmaceuticals and therapeutic medical device development, it has been estimated that only one in 5,000 to 10,000 innovations survive the long route from the initial findings to product commercialization [27, 28, 29]. This reflects a large gap between results from academic research and industrial product development. The different levels of product development are known as technology readiness levels, or TRLs (Table 1) [24]. Academic research generally ranges between TRL 1 to 3, while industry generally covers TRL 8 and 9. At TRL 4 to 7, the discovery process is often considered too applied for further scientific funding, whereas the industrial sector still considers it too risky to fund for market implementation.

**Table 1.** Technology Readiness Levels (TRLs) as instated by NASA [24].

| TRL | Definition |
| --- | --- |
| 1 | Basic principles observed and reported |
| 2 | Technology concept and/or application formulated |
| 3 | Analytical and experimental critical function and/or characteristic proof-of-concept |
| 4 | Component validation in laboratory environment |
| 5 | Component validation in relevant environment |
| 6 | System/subsystem model or prototype demonstration in relevant environment |
| 7 | System prototype demonstration in relevant environment |
| 8 | Actual system completed and qualified through test and demonstration |
| 9 | Actual system proven through successful operation |

Reasons for new technology to not survive this Valley of Death include cumbersome contracting or procurement of technology requirements, lack of exposure, lack of entrepreneurial management, lack of adequate funding for further development, and the lack of a strong link between technology development efforts and industrial deployment [27, 28, 29]. In an effort to decrease the amount of research that does not bridge the Valley of Death, funding agencies now often demand a direct collaboration between academia with industrial partners. Still, the Valley of Death remains as large as ever.

### Exploring the distance between academia and industry

To gain insights into this pressing matter, we conducted in-depth surveys amongst practitioners. To this end, we approached multiple industrial and academic experts to assess their perceptions on how to increase chances of research surviving the Valley of Death. Specifically, we aimed to ascertain I) what factors influence the choice for an industrial micro-organism and opportunities to close the distance between academia and industry.

Based on literature and experience [30, 31, 32, 33, 19, 34], a topic list was built to guide the interviews:

1. Choice of production platform was addressed by questions such as which production organism is used now, what factors may influence the choice for a production organism, including limitations or possible restrictions, company and set-up flexibility and what a production organism would need for the participating companies to switch.
2. Opportunities between industry and academia was addressed by questions including common ground, differences, challenges, opportunities, interesting fields of research, impact of science in industry, and collaboration. To this end, we included the presentation of a simplified overview of industrial production from the Design-BuildTest-Learn cycle [35] via laboratory scale to industrial production scale, where participants could indicate the main bottlenecks which indirectly represent the main short-term research opportunities for academia (Figure1). Figure 1 was used to stimulate in-depth discussion on challenges and opportunities, allowing us to construct a plan to navigate the Valley of Death.

**Figure 1.**
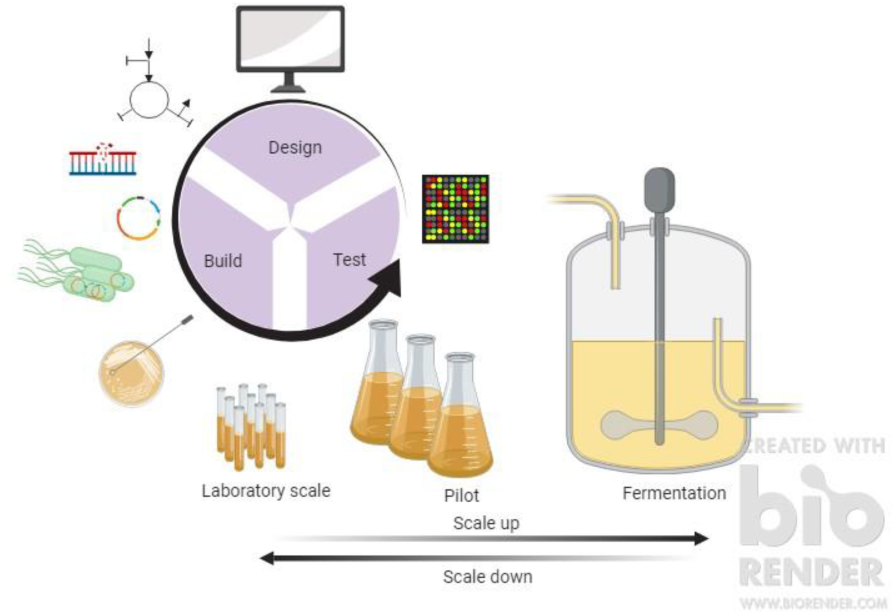
Overview of industrial development and production process. The schematic covers the main steps between the DBT-cycle and the industrial production scale.

## Methods

Because of a perceived lack of systemic exploration of the factors that influence if novel research makes it through the Valley of Death to market implementation, a qualitative and exploratory study was performed to gain a comprehensive insight into the perception of industry and academia by conducting interviews.

### Recruitment and Procedure

Recruitment took place by gathering a convenience sample based on our direct network and visited conferences. A list was devised of possible participants, from which 15 companies were selected based on their location, Europe or America, and field of research, pharmaceuticals, food, industrial chemicals, or production organism development. Some companies selected cover a combination of these fields. A variety of large and small and new and established companies were selected carefully from each continent, to determine the effect of a different regulation in industrial applications of bacteria and GMOs in particular, and over different companies from different sectors. Chief executive officers or chief technical officers were mainly approached to ensure the most overhead view on proceedings and the clearest perspective on the general aim, flexibility and workings of the company. In three cases, we were redirected to someone else within the company. Of the fifteen companies approached, nine agreed to participate, five did not answer, and one declined participation citing confidentiality over proprietary information. Of the nine companies that agreed, one decided to withdraw from participating as they could not obtain consent from their legal department after receiving the transcripts. The distribution of companies over continent and field are shown in Table 2.

**Table 2.**
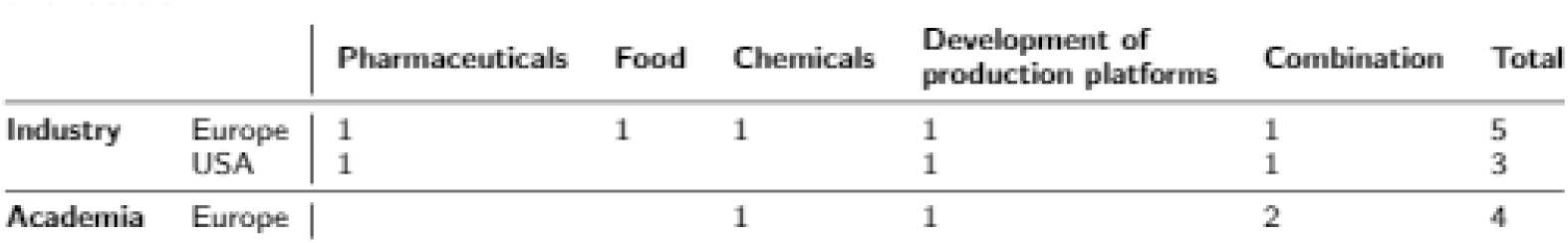
Participants in the interviews: the field occupied by participating companies and academics, and location.

|  |  | Pharmaceuticals | Food | Chemicals | Development of production platforms | Combination | Total |
| --- | --- | --- | --- | --- | --- | --- | --- |
| Industry | Europe | 1 | 1 | 1 | 1 | 1 | 5 |
|  | USA | 1 |  |  | 1 | 1 | 3 |
| Academia | Europe |  |  | 1 | 1 | 2 | 4 |

Of the convenience sample, 4 European universities were selected based on their diverting expertise, similarly to companies. All universities approached agreed to participate. Three of the academic participants had present or past experience in working at a company. All participants were engaged in collaborations with multiple industrial sectors. The distribution of companies over continent and field are shown in Table 2.

Potential participants were approached up to three times via email to make an appointment. If possible, interviews were conducted in person, otherwise face-to- face through Skype, WebEx or Bluejeans, according to the participants preference. During one interview, the camera of the participant failed and the interview was done by voice only. Both through e-mail and at the start of each interview the aim and methodology of the research was explained to the participants. Before the start of each interview, participants were presented with written information and contact details, and were asked to sign a declaration of consent. Interviews were conducted over a period of 8 months, from April to November 2019. After 8 interviews, the content of answers started to show overlap with previous interviews indicating data saturation.

### Interviews

Prior to data collection, the interview questions were discussed with an expert working in academia and working closely with industry, from which adjustments were made. Adjustments were focused on making the questions more open-ended and general to prevent bias and allow for a more in-depth discussion by leaving room for follow up questions (form S2). The use of appreciative inquiry in the questions [36] encouraged and inspired the participants to answer according to their own perspectives, ideas and experiences, as opposed to following a strict interview structure. In practice, this means that questions are asked with a positive spin, to focus more on solutions rather than on problems. A pilot interview with an expert working both in industry and academia was then held, indicating that the qualitative face-to-face interviews offered the expected in-depth information and clarity for both the participant and the interviewer, as it allowed for elaboration on specific topics, insight into technical differences or company aims. Special care was taken during the interviews to make sure there is a consensus of participants on definitions and terms used, by asking follow-up questions. Each interview was conducted by two members of the research team. The 13 interviews lasted on average 41 minutes, ranging from 26 to 54 minutes. With the participants’ verbal and written consent, all interviews were audio-taped and transcribed (intelligent verbatim style). The data was pseudonymized. When necessary, audio files were used to confirm transcripts and listen to excerpts within their original context. Transcripts and specific quotes used from each transcript were shared with the participants to consent upon prior to analysis. All participants responded, and minor editorial changes were made.

## Results

### Navigating the Valley of Death

Data was processed using the six steps for qualitative data analysis [37]. All analyses were done by the two interviewers, LK and EAG. In the first step, transcripts were made and carefully read. In the second step, codes were formulated manually using the research questions. Thirdly, the transcripts were coded using the formulated top-down codes, and where necessary deriving new codes from the data. The processing of the transcripts was conducted via inductive category formation using QCAmap [38], according to a thematic content analysis [39, 37]. The interviewers discussed their respective individual top-down and bottom-up codes to define a final set of codes. In the fourth step, all codes were clustered into the research questions: A) Choice of Production Platform, and B) Opportunities between Industry and Academia. In step five and six, recurring themes found to affect multiple codes were derived from the data and defined carefully. The nine themes found pertain either one or both research questions. The themes also led to the establishment of an overarching topic: all themes found relate to a difference in aims of industry and academia. To support the themes found, direct quotes are provided in the result section, selected to represent participants’ opinions, views and experiences. Table 3 presents an overview of the clusters, codes and derived themes. The main results of the analyses are presented below for each of the greater themes.

**Table 3.** Themes, research questions and codes of transcriptions. Top-down codes were formed before coding, bottom-up codes were derived during transcription.

| Themes | Top-down codes | Bottom-up codes |
| --- | --- | --- |
| Research Question 1: Choice of production platform |  |  |
| • I) Characteristics of microbial production platform<br>• II) Industrial process and set-up flexibility<br>• III) Regulations<br>• IV) Public perception | • What production platform is used<br>• Characteristics of used production platform<br>• Flexibility on industrial set-up | • Limitations on choice of production platform<br>• External influences on choice of production platform |
| Research Question 2: Opportunities between industry and academia |  |  |
| • V) Genome engineering<br>• VI) Scale-up<br>• VII) Increase strain robustness | • Opportunities and developments<br>• Distance between industry and academia<br>• Impact of industry on academia | • Personal or company aim<br>• Influence of industrial experiences |
| The aims of industry and academia |  |  |
| • VIII) Curiosity-driven versus solution-based research<br>• IX) The rise of small companies | • What production platform is used<br>• Characteristics of used production platform<br>• Flexibility on industrial set-up<br>• Opportunities and developments<br>• Distance between industry and academia<br>• Impact of industry on academia | • Limitations on choice of production platform<br>• External influences on choice of production platform<br>• Personal or company aim<br>• Influence of industrial experiences<br>• Communication |

#### (A) Choice of production platform

Four main themes pertain the choice for a certain production organism: I) Characteristics of microbial production platform, II) Industrial process and set-up flexibility, III) Regulations, and Public perception (Table 4).

**Table 4:**
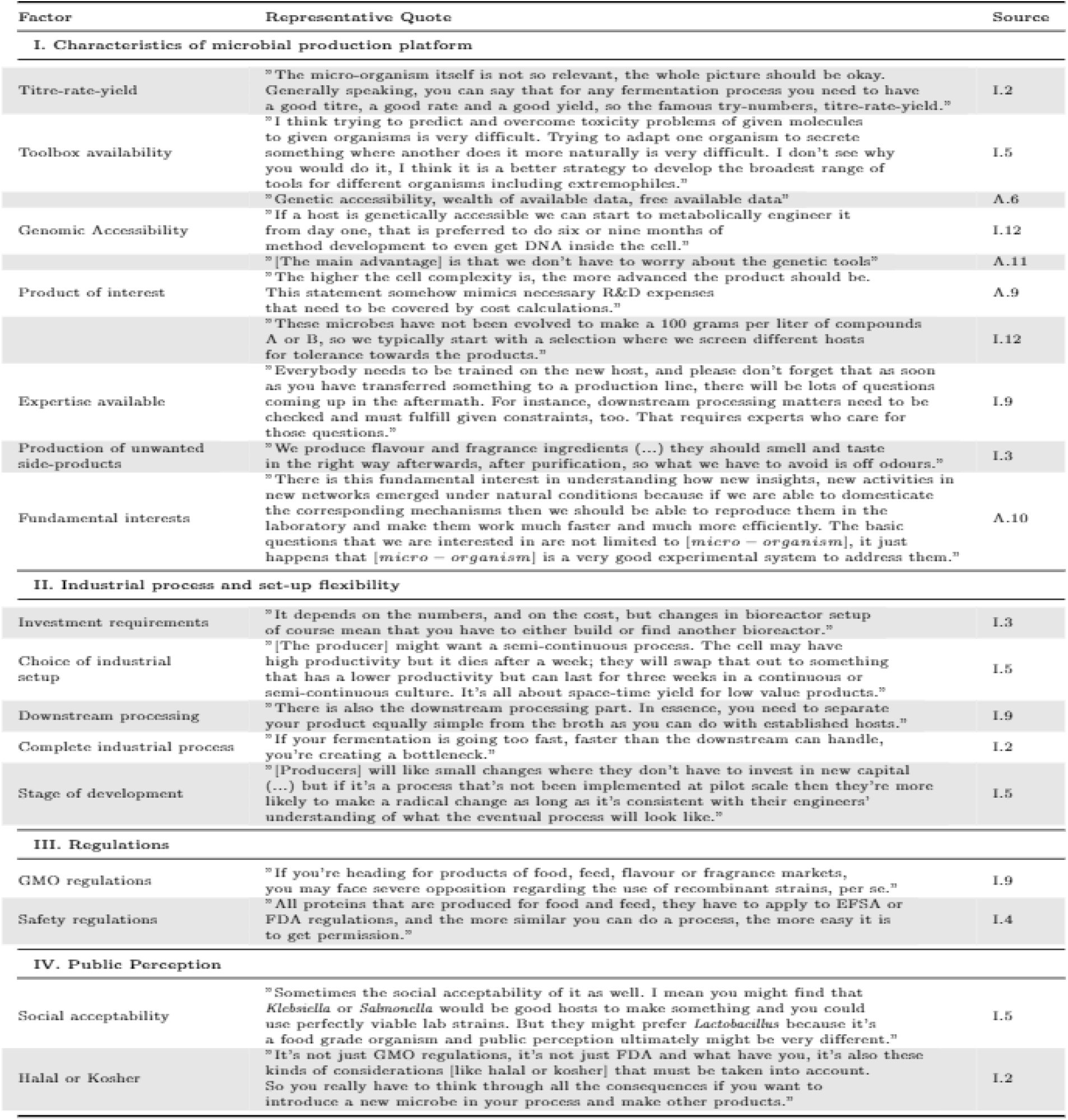
Factors affecting the choice of production platform with representational quotes from participants from industry (I) or academia (A), indicated by their numerical identifier.

| Factor | Representative Quote | Source |
| --- | --- | --- |
| **I. Characteristics of microbial production platform** |  |  |
| Titre-rate-yield | "The micro-organism itself is not so relevant, the whole picture should be okay. Generally speaking, you can say that for any fermentation process you need to have a good titre, a good rate and a good yield, so the famous try-numbers, titre-rate-yield." | I.2 |
| Toolbox availability | "I think trying to predict and overcome toxicity problems of given molecules to given organisms is very difficult. Trying to adapt one organism to secrete something where another does it more naturally is very difficult. I don't see why you would do it, I think it is a better strategy to develop the broadest range of tools for different organisms including extremophiles." | I.5 |
| Genomic Accessibility | "Genetic accessibility, wealth of available data, free available data" | A.6 |
|  | "If a host is genetically accessible we can start to metabolically engineer it from day one, that is preferred to do six or nine months of method development to even get DNA inside the cell." | I.12 |
|  | "[The main advantage] is that we don't have to worry about the genetic tools" | A.11 |
| Product of interest | "The higher the cell complexity is, the more advanced the product should be. This statement somehow mimics necessary R&D expenses that need to be covered by cost calculations." | A.9 |
|  | "These microbes have not been evolved to make a 100 grams per liter of compounds A or B, so we typically start with a selection where we screen different hosts for tolerance towards the products." | I.12 |
| Expertise available | "Everybody needs to be trained on the new host, and please don't forget that as soon as you have transferred something to a production line, there will be lots of questions coming up in the aftermath. For instance, downstream processing matters need to be checked and must fulfill given constraints, too. That requires experts who care for those questions." | I.9 |
| Production of unwanted side-products | "We produce flavour and fragrance ingredients (...) they should smell and taste in the right way afterwards, after purification, so what we have to avoid is off odours." | I.3 |
| Fundamental interests | "There is this fundamental interest in understanding how new insights, new activities in new networks emerged under natural conditions because if we are able to domesticate the corresponding mechanisms then we should be able to reproduce them in the laboratory and make them work much faster and much more efficiently. The basic questions that we are interested in are not limited to [micro – organism], it just happens that [micro – organism] is a very good experimental system to address them." | A.10 |
| **II. Industrial process and set-up flexibility** |  |  |
| Investment requirements | "It depends on the numbers, and on the cost, but changes in bioreactor setup of course mean that you have to either build or find another bioreactor." | I.3 |
| Choice of industrial setup | "[The producer] might want a semi-continuous process. The cell may have high productivity but it dies after a week; they will swap that out to something that has a lower productivity but can last for three weeks in a continuous or semi-continuous culture. It's all about space-time yield for low value products." | I.5 |
| Downstream processing | "There is also the downstream processing part. In essence, you need to separate your product equally simple from the broth as you can do with established hosts." | I.9 |
| Complete industrial process | "If your fermentation is going too fast, faster than the downstream can handle, you're creating a bottleneck." | I.2 |
| Stage of development | "[Producers] will like small changes where they don't have to invest in new capital (...) but if it's a process that's not been implemented at pilot scale then they're more likely to make a radical change as long as it's consistent with their engineers' understanding of what the eventual process will look like." | I.5 |
| **III. Regulations** |  |  |
| GMO regulations | "If you're heading for products of food, feed, flavour or fragrance markets, you may face severe opposition regarding the use of recombinant strains, per se." | I.9 |
| Safety regulations | "All proteins that are produced for food and feed, they have to apply to EFSA or FDA regulations, and the more similar you can do a process, the more easy it is to get permission." | I.4 |
| **IV. Public Perception** |  |  |
| Social acceptability | "Sometimes the social acceptability of it as well. I mean you might find that *Klebsiella* or *Salmonella* would be good hosts to make something and you could use perfectly viable lab strains. But they might prefer *Lactobacillus* because it's a food grade organism and public perception ultimately might be very different." | I.5 |
| Halal or Kosher | "It's not just GMO regulations, it's not just FDA and what have you, it's also these kinds of considerations [like halal or kosher] that must be taken into account. So you really have to think through all the consequences if you want to introduce a new microbe in your process and make other products." | I.2 |

The factors that influence the choice of production platform in industry and academia can be divided into technical factors, such as the characteristics of the production platform and industrial process and set-up flexibility, sector-specific factors like regulations, and social factors like public perception. In industry are many technical, sector-based and social restrictions to choosing a production host. As they must keep the end-user in mind throughout the entire development and production process, products must be made as cost-effective, fast, environmentally-friendly and safe as possible.

For academia, mostly technical factors play a role when deciding to work with a specific micro-organism. These boil down to availability of a suitable genetic and biochemical toolbox, genomic accessibility, product of interest and fundamental interests. Only the latter is specific to academia, the rest is shared with industry. This means that academics are more free to work with any micro-organism, even with microbes that cannot be applied (yet) to industrial processes.

#### (B) Opportunities between Academia and Industry

To determine what the main future opportunities are, participants were asked for future research of interest, the main bottlenecks in their production process and possibilities, and what current scientific progress they expected to have most future impact (Figure 2).

**Figure 2.**
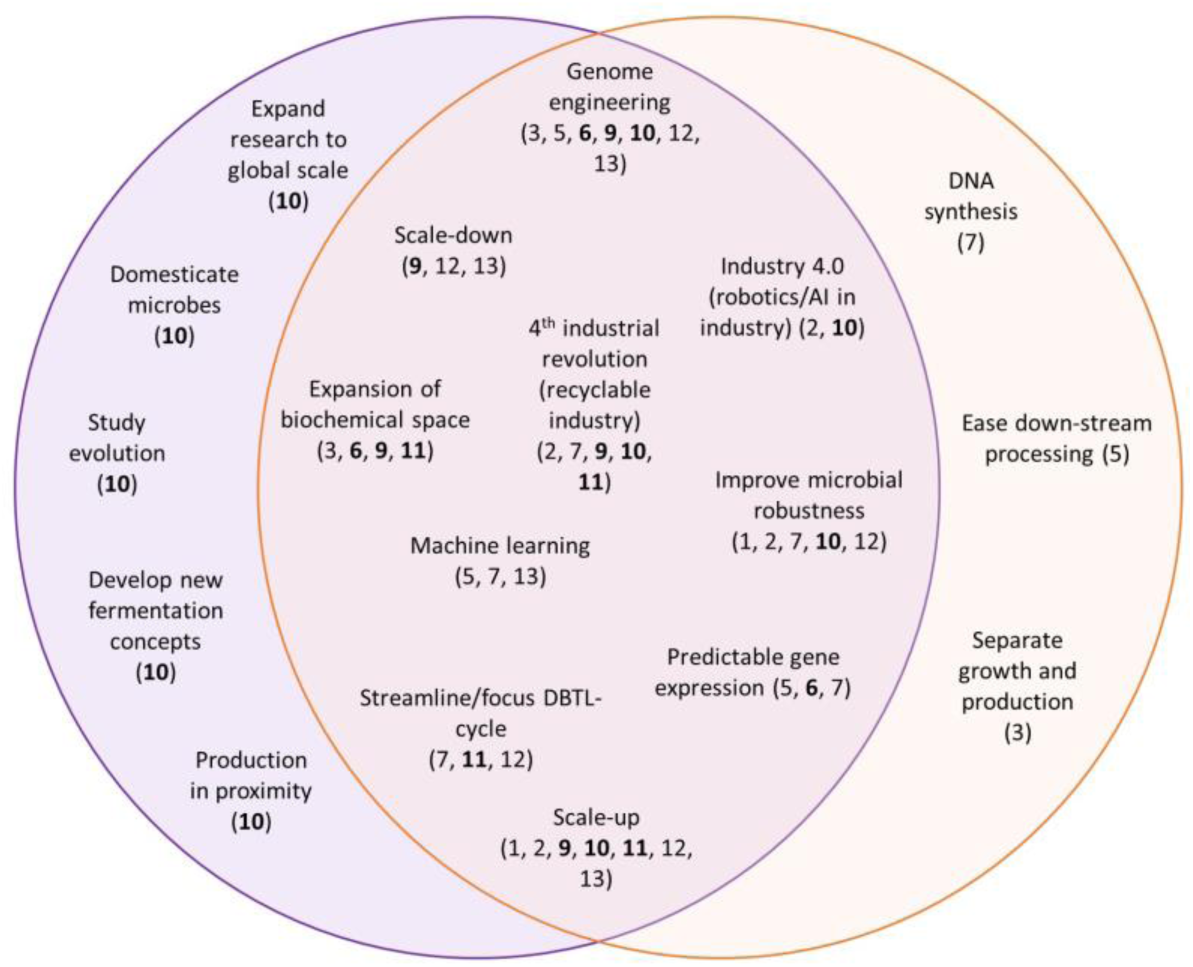
Venn diagram of research areas of interest per field. A schematic overview of the research areas indicated as interesting by academia (left, purple) and industry (right, orange). Overlapping research of interest is in the middle. Per subject, the identifier of each participant is indicated. Identifiers of academic collaborators are printed in bold.

From this overview, three main technical themes emerged: V) Genome engineering, VI) Scaleup, and VII) Improved robustness. These will be discussed in more detail below.

#### V) Genome engineering

When presented with Figure 1, most industrial participants identified the whole or part of the DBTL-cycle as the main opportunity for improvement.

“You test [a new design], you find something out and then you have to design, you get a lot of data, you get some results, you don’t understand them, and you have to analyse. The learn part is the bottleneck because all the other things are automatable or scalable to some extent, so the learning bit is not.” Interviewee 3

For industry, the possibility to affordably construct an entire genome combined with the rise of novel tools such as CRISPR-Cas for genome engineering significantly ease this bottleneck.

“On the build side [of the DBTL-cycle] there’s a lot of change. Synthetic DNA is becoming cheaper and cheaper by the day, sequencing is becoming cheaper by the day, CRISPR is like 6 years old and [recently] there was a new version released that really put the whole town plus all of the genetic engineering upside down. [...] It’s just changing the timelines and the ability so much.” Interviewee 12

For academia, such tools pose promising ways to improve upon fundamental research, which is required for understanding strain design.

“I would hope that it will accelerate the way at which we can generate industrially meaningful strains, and it will expand the extent of manipulation that we can achieve. So rather than looking at one gene or one pathway, we should hopefully be able to look at one entire section of metabolism in one go, and this would hopefully have a very good influence on the strain design that we want to achieve.” Interviewee 6

#### VI) Scale-up

To reliably scale-up results obtained in perfect lab-conditions to industrial scale is challenging at best. Scaling-up was recognised as the second main bottleneck. Even in academia, the importance of reliable scale-up to bridge the Valley of Death was recognised:

“You can develop the most sophisticated CRISPR system, the most sophisticated recombineering, the most sophisticated reactions, you can do wonderful things in the lab: if you cannot scale them up, then industry will not be interested, period.” Interviewee 10

From an industrial point of view, there are few different reasons why scale-up can be a bottleneck, besides technical difficulties:

“The availability of equipment is actually the whole bottleneck. There is not a lot of pilot facilities in the US, so either you build your own or you go to Europe for piloting. That’s non-trivial. I think there’s a lot of syn-bio companies that are mostly comprised of genetic engineers or metabolic engineers that have never seen a large-scale fermentor and a large scale production process including purification. So if you start a scale-up trajectory without people that know what they’re doing then it’s going to be a bottleneck. But if you have a team, [...] then you should be able to do that in a pretty smooth trajectory. There’s always going to be surprises and hiccups. What I’ve also seen is that lots of syn-bio companies have very aggressive timelines, probably due to their investors pushing really hard, and then they are going to take short-cuts and that’s going to blow up in your face. There is not a technical bottleneck, but it’s more of a managerial bottleneck at that point.” Interviewee 12

#### VII) Improved strain robustness

When asked directly, all participants indicated that the robustness of their used or preferred production platform could be improved. Reasons for improving robustness circle back to scale-up without fail, since industrially used production organisms are grown in conditions that are strictly controlled.

“Robustness in an industrial sense is that it is operationally robust: meaning it is not very sensitive to infections, it is not very sensitive to minor variations that you always have in the conditions of the process, e.g. due to scale. So there should be some tolerance in the microbes to slight variations in the process conditions, which of course we try to control that as much as possible.” Interviewee 2

In the few cases in which robustness was not recognised as an issue, it was solely because the production platform of interest already performed optimally at large scale:

“We experience our micro-organism is very robust under the conditions that are relevant to us, that is industrial fermentors, so we don’t see a need for further improvement there.” Interviewee 3

Robustness is thus only deemed important to aid production platforms to perform reliably on industrial scale. Even in academia, this link was made.

“Fluctuations on large scale are much more prominent then on small scale, and constant performance through larger fluctuations in large scale most probably could be improved.” Interviewee 6

Overall, the long- and short-term interests of academia and industry align very well. However, there were some differences in interest. In industry, interests are often very practical, all leading to an increase in titre-rate-yield. Subjects included the use of a different C-source, separated growthand production cycles, and ease down-stream processing. Amongst academics, topics of interest that did not align with those in industry were less aimed at application and practicality, but of a more fundamental or futuristic nature. These included for example the development of new fermentation concepts, expanding of research to global scale and studying evolution.

### The aims of industry and academia

From the in-depth interviews, an overarching topic emerged. All nine themes together lead to a difference in aim between industry and academia. Two themes where found vital in this difference in aim, and as such influence both research questions: VIII) Curiosity-driven versus solution-based research, and IX) the rise of small companies.

#### VIII) Curiosity-driven versus solution-based research

The precarious balance between curiosity-driven and solution-based research, the latter mainly with industrial application in mind or on stakeholders’ demand, can be defined most clearly by looking at the collaborations between academia and industry. All participants, both from industry and academia, indicated current involvement in collaborative projects. However, these collaborations are not necessarily initiated by industry nor based on industrial interests. By academics, it was more described as follows:

“It’s rarely the case that somebody [from industry] comes to me and says “This is something that you should work on”, but of course you talk to people at conferences, at seminars, and you try to understand what the problems are, and you try to find out if you can solve some of these problems with the tools that you have available. But it’s a bit of a balance, I try to make sure that a substantial part of our activities does not have a direct line. In a way, it’s curiosity driven.” Interviewee 6

Collaborations between academia and industry are not based on a demand-supply model, but rather on academics trying to fill existing or predicted gaps in industry. All academic partners did indicate a strong influence on their work from industry, even if not directly. However, a way to deal with this potential struggle was explained:

“We are driven by curiosity and obviously we have to frame our curiosity and our personal interest in the interest of our sponsors, and in the interest of society, and this is perfectly fine. Just remember that the big artist in the Renaissance were sponsored by the popes, by the emperors and so, and [the artists] had to produce things that [the sponsors] liked. I think this is not completely shocking that we are constrained by the ideas of our sponsors. I think that our challenge [...] has a point even of enjoyment. It is how you connect your own personal interest and our own curiosity with these big demands that your sponsors have for your research.” Interviewee 10

Industry thus does not approach academia with questions, but rather shops around for interesting opportunities they can easily adapt and apply. This indicates that they rely heavily on research performed in academia to make large strides in innovation.

Industry indicates this when finding new investment opportunities:

“The biggest opportunities lie in the combination for a company like us, or a hospital research group for example, that have developed something novel, something proprietary, but [they, academia] have no understanding of the hurdles they’ll face or the means to demonstrate its viability to be a competitive product in the market. Because that’s what’s going to be necessary to attract investment. To give it a chance of survival.” Interviewee 5

The clear distinction between technical, sector-driven and social factors influencing main research choices and definition of opportunities can be derived from the difference in the main aim of each field. Where industry looks for applications and adaptable solutions to day-to-day problems, academics aim to discover answers to fundamental questions without being limited by end-user demands. Communication to align these different aims towards one goal is vital:

“First of all, there has to be good communication, because academia works in a completely different way to industry. Industry is all about timelines, deadlines, deliverables, project management, progress towards measurable output. Academia is about testing, discovery, interesting products, interesting processes, new methods, means of discovering where science can go.” Interviewee

#### IX) The rise of small companies

Small companies that arise from academia to commercialise promising academic findings are called start-ups. Start-ups do not only focus on the academically recognised technical factors, but include the sector-based and social factors that might influence the successful industrial introduction of their product, focusing specifically on TRL 4-7.

Some see start-ups as a natural way to close the gap between academia and industry, by presenting interesting research ready for commercialisation on a silver platter:

“It’s those partnerships in between, it’s the networking and the identification of viable product opportunities that are emerging from either academic groups, hospital groups, virtual companies and then the partnerships that takes that feasibility demonstration onto viability. That’s where the opportunity is. Bridging that gap from feasibility to viability. Because that’s what derisks it, and brings in funding to take it forward. I think otherwise so many things can sit in academia, very interesting, very exciting, and probably more so in Europe than in the United States, and they just don’t see the light of day because they haven’t taken the next step and that’s where certainly a company like ours, but other small companies can come in and make that difference.” Interviewee 5

Other interviewees indicate that the rise of start-ups widens the communicative and collaborative distance between academia and industry. However, they do understand why companies prefer dealing with start-up companies over academics:

“What we observe is that all these new developments, new technologies and everything, is mostly the business of small companies. The big companies just become very conservative. [...] Big companies prefer to deal with smaller companies than dealing directly with scientists, and the reason [...] is that scientists have too big a mouth. That means that the moment you have discovered something interesting you immediately start spreading the word, and this is something that the companies don’t like at all.” Interviewee 10

## Discussion

### The importance of technical, sector-based and social factors

Three groups of factors were recognised when considering the choice of microorganisms currently applied in industrial biotechnology: technical, sector-based and social. Current host systems widely applied include bacteria, yeasts, filamentous fungi and unicellular algae. Industrially applied strains all come with their own strengths and drawbacks [40, 5, 41]. Technical factors that influence the choice for production platform comprise, among others, the ability to produce complex chemicals, growth rate, the ability to perform post-translational modifications and the use of cheap medium components. Sector-based factors include the used micro-organism to be a natural producer of the product of interest, the cultivation and production process to fit within the complete industrial process, or familiarity with working with a specific strain from out the company history. Social factors for the wide application of these production hosts include public familiarity and the safety status of a species.

Academic participants only cited technical reasons for the selection of a microorganism such as ease to work with, genomic accessibility and wealth of publicly available data. In industry, sector-based factors such as product of interest, and social influences derived from demands of the end-user or regulations applied by the government also strongly influence choices made. Still, the main research opportunities and future prospects found are of a technical nature. Genome engineering opportunities, predictability, stability, scale-up, and increased strain robustness all focus on improving the production process from a technical point of view. Mismatches leading to an innovation not making it through the Valley of Death thus may occur when the social and sector-based aspects are not kept in mind.

Changing the production organism used in an established industrial process is considered a stark industrial shift, which heavily depends on research. Adapting an existing production process is expensive, requires specific scientific knowledge and takes time to develop and test. As changing a production platform is generally bound to affect the industrial setup, the gain in yield and characteristics has to be substantial to make up for the cost of the investment on set-up changes. Additionally, microbes with possibly more interesting traits to better produce specific substances are being discovered every day. It is thus not surprising that there is a lot of research focused not on finding the perfect production host for specific purposes, but on the de novo design and construct of one. This is either done by making a cell including only traits specifically selected as required for survival and production, or by re-wiring and streamlining an existing production microbial platform.

Computational methods are paving the way towards big data processing towards one goal of interest, in any available species. However, implementation and testing of many different designs through the DBTL cycle and predictability of gene expression were identified as the bottlenecks towards progress, by both industrial and academic participants.

### The rise of start-ups

The main reason for the distance between academia and industry arises from the difference in perspective. Where academia focuses on understanding a microorganism, industry always focuses on the entire production process. At the same time, academia is encouraged to perform curiosity-driven research, answering fundamental questions. Where industry works on solving the problems of today, academia focuses on the problems of tomorrow. In spite of this difference in aim, both parties need each other for funding, direction, creativity, innovation and exploring boundaries. Both parties also agree that communication could be improved to improve cooperation and increase opportunities for process or product development and introduction to the market.

These differences found between the general aims, way of thinking and factors included in decisionmaking between academia and industry can be largely brought back to the different technology readiness levels (TRLs) they operate on [24]. As earlier indicated, academia works mostly at TRL 1-3 while industry tends to focus on TRL 8-9. The enormous gap left from TRL 4-7 is the most high-risk phase in the development of a new product or process.

To close this gap, a solution may have naturally emerged: start-up companies. Rather than direct translation of academic research in industry, new companies emerge straight out of academia, which are often acquired by large companies when proving particularly promising. Start-up companies are closely linked to both industry and academia, allowing for a better insight in the restrictions and aims of either. They are set up specifically to bridge TRL 4-7, more willing to take risks, and can change their production platform more easily and quickly if so needed. The continuous process of using novel advanced technologies to build a firm has moved to the heart of innovation strategy [42]. Start-up initiatives focusing on the technical, sector-based and social factors by presenting the best, most lucrative results achieved in academia on a silver platter, attracting their own initial funding and stirring commercial interest. This trend is currently seen throughout many different sectors, with the well-known example of the once-startup multinational IT company Facebook acquiring multiple startups [43].

In the early twentieth century, government-funded science fostered a first academic revolution. Entrepreneurial science, where collaborations were formed with industry, chimed in the second academic revolution [44]. Nowadays, government funded research often even requires collaborations with industrial partners, as seen in European projects such as EmPowerPutida, P4SB or IBISBA [45, 46]. The swift rise of start-ups has chimed in a third academic revolution, where securing the IP of novel discoveries will grow in importance. This has impacted the role of academia, which is pushed towards faster development of more applied research.

All academic and some industrial participants have indicated a growing distance between academia and industry, impacting the communication and collaborations. Some have indicated this growing distance to be worrisome. As start-up companies present commercialized techniques that can be quickly integrated in any company, they mask the years of university-driven research and development that occurred before forming the start-up. Industry then much rather obtains the techniques or products from the start-up company directly than investing in the academic research.

However, the natural rise of start-ups should be seen as an opportunity. Startups can ease the transition of academic research to actual industrial application, closing the Valley of Death by forming an independent bridge between academia and industry. It also opens up room for academics to focus on what they do best: education and research. It was earlier recognised [47] that having opportunities in academia to translate their research effectively into concrete products benefits academic institutions, faculty members, industry, and society [44]. A case could be made for collaborations to include some attention for the possibility of development of start-ups, which might lead to more attention to promising results and how to grow them into market applications from an early point on.

### Strengths and limitations of the study

After approximately 8 interviews data saturation occurred. This means that no new information was generated and hence indicates a good reflection of the target community. The large spread of different companies over different biotechnology sectors, of different sizes and spread over different locations around the globe allowed to reasonably generalise the findings so that the universal bottlenecks, challenges and opportunities could be easily recognised. However, another interesting approach to this research could be in the subtle differences from companies in different continents. A larger sample would be more reflective of the continental community. Companies and research facilities from other continents could add to this. Additionally, only European academic instances were included in the research.

## Conclusions

Within industrial biotechnology, research on a suitable production platform is paramount. Existing production platforms are improved, or new ones are developed. From an academic point of view, accessibility and ease to work with are the main reasons to choose for one micro-organism. For industry, technical factors such as the intrinsic characteristics of the production platform, industrial process and set-up flexibility, sector-based factors such as regulations and social factors such as public perception are important influences. To ensure research innovation makes it through the Valley of Death to market commercialisation, these factors must be taken into account early on. Universally recognised research topics of interests are genome engineering, scale-up, and improved strain robustness. The main cause for lack of market introduction are the vastly different technology readiness levels that academia and industry operate on. Industry always focuses on solution- based research to improve the titre-rate-yield, while academia focuses on curiosity- driven, fundamental research, steps removed from industrial application. Neither party wants this to change, while both indicate a need for improved communication so that no opportunities are lost. Start-ups can serve as the bridge over the Valley of Death, connecting feasibility to viability, by acting as a communication channel between academia where the research originates, and industry to help set-up and fund along the way.

## Author’s contributions

Conceived the in silico study: VAPMdS/LFCK Literature research: LFCK Interview setup: LFCK/EAG/AW Interviews: LFCK/EAG Data Analysis: LFCK/EAG/AW Work Supervision: AW/PJS/VAPMdS Wrote Manuscript: LFCK Proofread Manuscript: AW/PJS/VAPMdS Arranged Funding: VAPMdS

## Acknowledgements

We thank all our interviewees for participating in our research. We thank dr. R. A. Weusthuis for his advice on the interview.

